# Evolutionary diversification of tiny ocean predators

**DOI:** 10.1101/2020.10.09.333062

**Authors:** Francisco Latorre, Ina M. Deutschmann, Aurelie Labarre, Aleix Obiol, Anders Krabberød, Eric Pelletier, Michael E. Sieracki, Corinne Cruaud, Olivier Jaillon, Ramon Massana, Ramiro Logares

## Abstract

Unicellular eukaryotic predators have a crucial role in the functioning of the ocean ecosystem by recycling nutrients and energy that are channeled to upper trophic levels. Traditionally, these evolutionary-diverse organisms have been combined into a single functional group (Heterotrophic flagellates), overlooking their organismal differences. Here we investigate four evolutionary related species belonging to one cosmopolitan family of uncultured marine picoeukaryotic predators: MAST-4 (species A, B, C, and E). Co-occurrence and distribution analyses in the global surface ocean indicated contrasting patterns in MAST-4A & C, suggesting adaptation to different temperatures. We then investigated whether these spatial distribution patterns were mirrored by MAST-4 genomic content using Single-Cell Genomics. Analyses of 69 single-cells recovered 66-83% of the MAST-4A/B/C/E genomes, which displayed substantial inter-species divergence. MAST-4 genomes were similar in terms of broad gene functional categories, but they differed in enzymes of ecological relevance, such as glycoside hydrolases (GHs), which are part of the food degradation machinery in MAST-4. Interestingly, MAST-4 species featuring a similar GH composition co-excluded each other (A & C) in the surface global ocean, while species with a different set of GHs appeared to be able to co-exist (species B & C) suggesting further niche diversification associated to prey digestion. We propose that differential niche adaptation to temperature and prey type has promoted adaptive evolutionary diversification in MAST-4. Altogether, we show that minute ocean predators from the same family may have different biogeography and genomic content, which need to be accounted to better comprehend marine food webs.

## INTRODUCTION

Ocean microbes are fundamental for the functioning of the Earth ecosystem, playing prominent roles in the global cycling of carbon and nutrients (1). In particular, small phototrophic microbes are responsible for ~50% of the primary production on the planet (2). In turn, heterotrophic microbes have a fundamental role in nutrient cycling and food-web dynamics (3). Heterotrophic flagellates, along with marine viruses, maintain prokaryotic and eukaryotic picoplankton at relatively stable abundances (4). At the same time, they transfer the organic matter they consume from lower to upper trophic levels, thus being a key component at the base of the ocean’s food web.

Among heterotrophic flagellates, Marine Stramenopiles (MASTs) play a prominent role in unicellular trophic interactions in the global ocean (5). MASTs are polyphyletic, including so far 18 subgroups (6). Except for a handful strains, MASTs remain uncultured (7), which complicates the study of their cell physiology, ecology and genomics. Studies using FISH (8–10) and metabarcoding (5, 11) helped to determine MAST cell sizes (2-5 μm), vertical and horizontal distributions in the ocean as well as metabolic activity. Further studies linked MAST’s cell morphology with environmental heterogeneity, for example, MAST-1B cell size varies with temperature (9). Other studies provided insights on the predatory behaviors of some MAST species. For instance, MAST-4 prey on *Synechococcus* (5) and SAR11 (12), two of the most abundant microorganisms in the ocean (13, 14). MAST-4 is a prominent clade within the MASTs, featuring small cells (2-3 μm), high relative abundance in comparison to other heterotrophic flagellates, and worldwide distributions (15). Due to these characteristics, MAST-4 can be considered as a model heterotrophic flagellate.

MAST-4 is constituted by at least 6 recognized species: MAST-4A/B/C/D/E/F based on 18S rRNA gene phylogenies (6). The biogeography of specific MAST-4 species has been partially elucidated: MAST-4 A, B and C occur in temperate and warm waters (17 – 30 °C), whereas species E prefers colder waters (2 – 17 °C) (16, 17). This suggests that MAST-4 species have adapted to different niche temperatures. MAST-4 biogeography could also be controlled by bottom-up or top-down biotic factors, such as prey/food-availability (e.g. bacteria, algae, Dissolved Organic Carbon) or predation respectively. A number of studies pointed to a positive correlation between prokaryotic and heterotrophic flagellates’ abundances (18–21); yet, it is unclear to what extent such biotic relationships can generate biogeography in MAST-4.

Biogeographic patterns of MAST-4 species can provide insights on the drivers that have promoted their evolutionary diversification. Identifying species-specific gene-functions, genes or gene variants may point to differential adaptations conferring higher fitness in specific biotic or abiotic conditions. In a heterotrophic flagellate model system like MAST-4, featuring a bacterivorous lifestyle, a first approach for assessing species-specific adaptations is to analyze Ecologically Relevant Genes (ERGs), which are those that could reflect associations with environmental heterogeneity or different ecological roles. Candidate ERGs are the enzymes present in the lysosome that are involved in phagocytosis, allowing the degradation of a wide variety of substances such as proteins, carbohydrates or nucleic acids among others (22). In heterotrophic flagellates, lysosomal enzymes are of particular relevance because different suites could potentially be associated to the degradation of different food items. Among them, Glycoside Hydrolases (GHs), commonly found in lysosomes, catalyze the hydrolysis of glycosidic bonds in complex sugars allowing the cell to digest other organisms and eventually feed on them. For example, the lysozyme (N-acetylmuramide glycanhydrolase) is a well-known enzyme under the GH category that catalyzes the breaking of the peptidoglycan cell wall found in bacteria (23). Other studies have shown that each MAST lineage may have a different functional profile in terms of organic matter processing (16).

Genomes are key to obtain ERGs from a species; common protocols require thousands if not millions of cells for genome sequencing. In uncultured protists such as MAST-4, obtaining this number of cells without a culture is practically an impossible task. This issue is circumvented with Single-Cell Genomics (SCG) (13, 24). The principles of this method consist in isolating single cells using flow cytometry, lysing the cells and amplifying and sequencing their genomes producing Single Amplified Genomes (SAGs). In previous work, Single-cell genomics allowed recovering ~20% of the MAST-4 genomes from individual cells, which increased to ~80% when genomes from different cells were co-assembled (16, 25). Here, we use the SAG collection produced by the *TARA Oceans* expedition (26), which generated 900 SAGs from 8 stations in the Indian Ocean and the Mediterranean Sea. We compiled the largest collection to date of MAST-4 SAGs, totaling 69 SAGs (23 MAST-4A, 9 MAST-4B, 20 MAST-4C and 17 MAST-4E). Using this novel dataset, together with other large metaomics datasets (metabarcoding, metagenomics and metatranscriptomics) from the *TARA Oceans* and *Malaspina-2010* expeditions (27) we address the main following questions: How different are MAST-4 species at the genome level? Did MAST-4 species diverge via niche adaptation? If so, is such adaptation reflected in their genomes and potential ecological interactions? Can ERG composition and expression provide insights on MAST-4 niche diversification?

## RESULTS

### MAST-4 global distributions and associations

MAST-4A/B/C/E Operational Taxonomic Units (OTUs; “species” proxies) tended to display specific spatial distributions in the global ocean, in some cases markedly contrasting (**Figure 1**). Specifically, species A and C were abundant and widespread across the global ocean and even though both may appear in the same sample, they tended to exclude each other, as indicated by their association sign (**Figure 1**). For example, in the Pacific Ocean when moving from equatorial waters to the north, there was a partial replacement between MAST-4C and A (See arrows in **Figure 1**). Species B displayed a more restricted distribution and a lower abundance when compared to species C and A, being more prevalent in the tropical and subtropical Atlantic Ocean (**Figure 1**). Our analyses indicated that species B co-occurred with species C, with both species co-excluding from species A (**Figure 1**). Species E had a lower abundance than the other species in the tropical and subtropical global ocean, with a distribution being limited to a few locations, mostly coastal areas (**Figure 1**). Species E had a weak negative association with MAST-4B (**Figure 1**).

**Figure 1.**
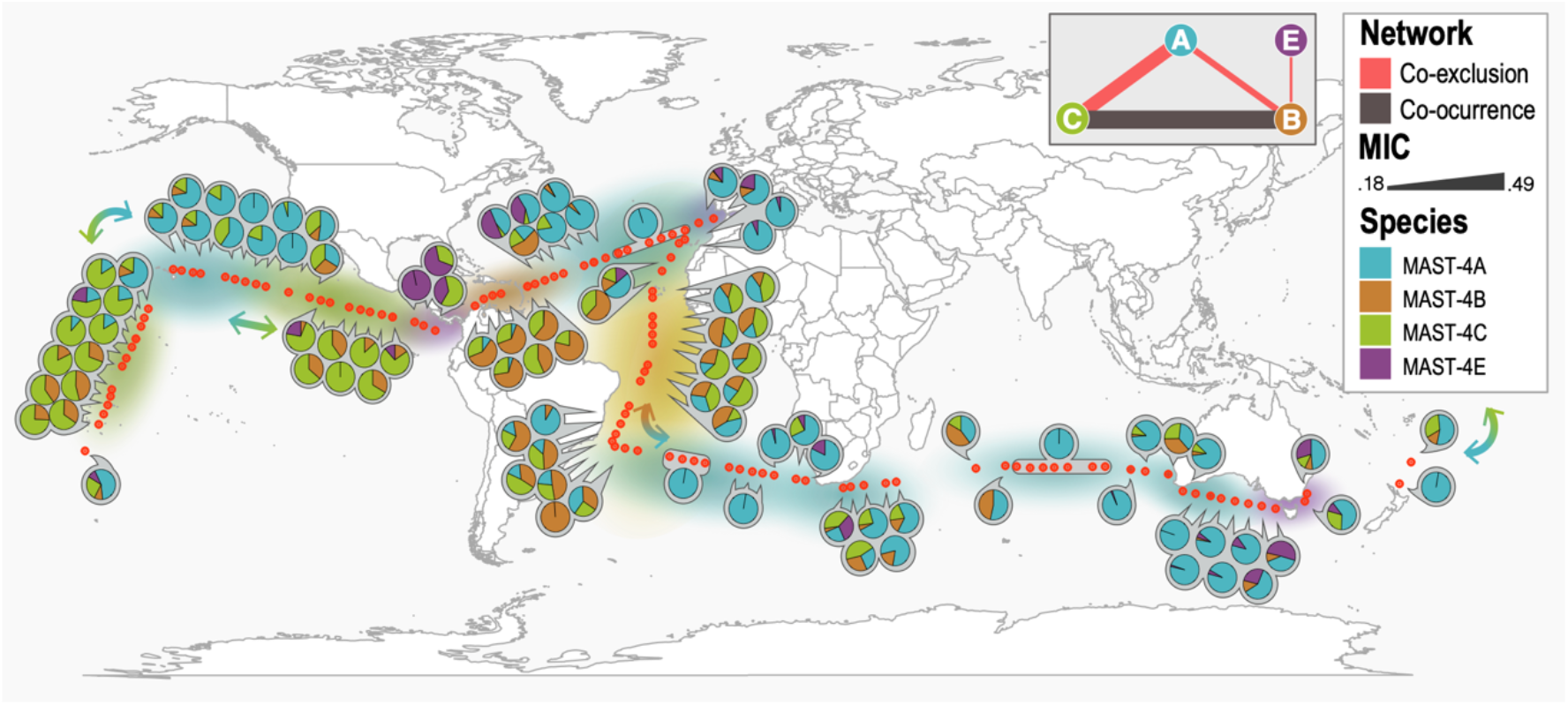
Distribution of MAST-4A/B/C/E species in the surface global ocean as inferred by OTUs based on the 18S rRNA gene (V4 region). Red dots show Malaspina stations while pie charts indicate the relative abundance of MAST-4 species at each station. The top-right inset network shows the association patterns between each MAST-4 species as measured using MIC analyses. The width of the edges in the network shows association strength as indicated in the legend (MIC). Background color shows the most abundant MAST-4 species in the region. Arrows point to areas with an important switch of the abundant species: note that the most abundant species, A and C, alternate predominance in large oceanic regions.

We have also investigated the association patterns between MAST-4A/B/C/E OTUs with other picoeukaryotes and prokaryotes. We found a total of 258 associations with other picoeukaryotic and 18 with prokaryotic OTUs that cannot be explained by the measured environmental factors (**Figure 2A**). MAST-4C and MAST-4B displayed the largest number of associated OTUs, 191 and 174 respectively, while MAST-4A, despite being abundant and cosmopolitan, had only 23 associations. MAST-4E had only 3 associations to other taxa different from MAST-4 (**Figure 2A**). Most associated taxa were related to a unique (59.3%), or to two (38.9%) MAST-4 species (mostly species B and C) [**Figure 2A**]. The co-occurring species B and C displayed the largest number of shared associated taxa (total 98 taxa), which in most cases (97%) were positively associated (**Figure 2A**). A fewer number of associations (total 13) was shared by the mutually excluding species A and C and, as expected, had opposite signs (50% positive and 50% negative; OTUs positively associated to MAST-4A were negatively associated to MAST-4C and vice versa) [**Figure 2A**]. A similar trend was observed between OTUs associated to species A and B (**Figure 2A**).

**Figure 2.**
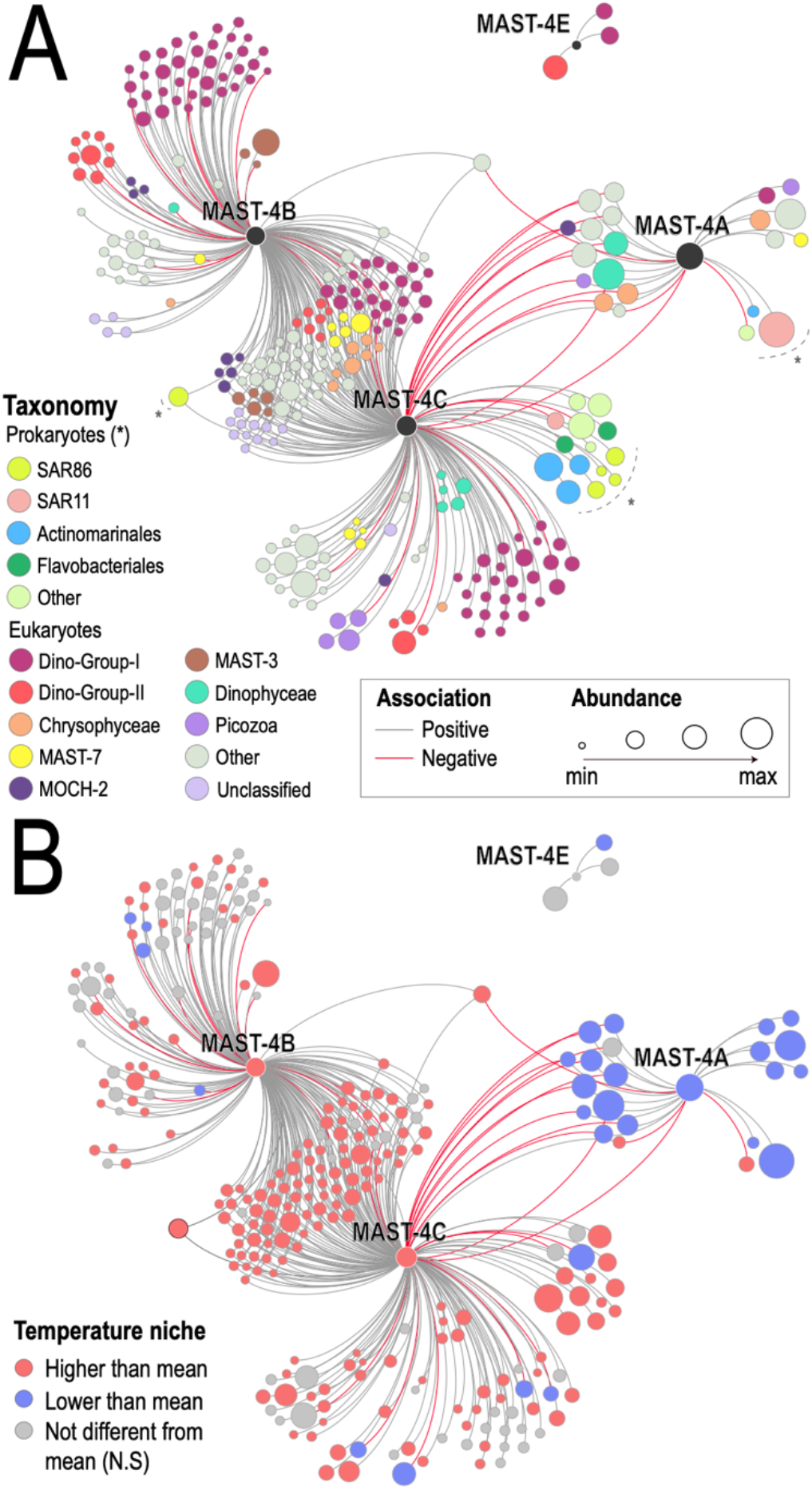
Association network including MAST-4 species, associated prokaryotes, and other pico-eukaryotes from the Malaspina expedition. Only OTUs with abundances >100 reads and occurrences >15% of the stations were considered in MIC analyses. A filtering strategy was applied to remove indirect (i.e. environmentally-driven) and weak associations (see Methods). Node size is proportional to the clr transformed abundance sum (see Methods). In **Panel A** nodes are colored based on taxonomy while in **Panel B** node color indicates whether specific OTUs displayed weighted mean temperatures significantly lower or higher than the unweighted mean temperature (~24.5 °C), pointing to species with temperature distributions that differ from chance. Note that MAST-4A and both MAST-B/C tend to show co-occurrences with other OTUs that display coherent temperature preferences. * N.S – Not Significant.

The most represented eukaryotic classes in the network included parasites (Syndiniales; 40.7% of the OTUs) and other marine Stramenopiles (16.8%), including MAST-1/3/7/11/25 and other MAST-4 OTUs belonging to species B/C/E, which had different 18S-V4 sequences when compared to those from the SAGs. The most represented prokaryotic classes in the network included the heterotrophic species SAR86 (1.8%) and the small-sized marine Actinobacteria (Actinomarinales; 1.4%) (**Figure 2A**). Other ecologically relevant classes that were present but displayed fewer OTUs were the eukaryote Picozoa (2.14%), which have similar physiological characteristics to MAST-4 (28, 29), or the prokaryotic SAR11 (0.71%), one of the most abundant bacteria in the ocean (14).

We analyzed the niche preference of individual MAST-4 OTUs as well as that of associated OTUs from other taxa in terms of temperature, salinity, NO_2_, NO_3_, PO_4_, SiO_4_ and fluorescence (**Supplementary table S5**). Adaptation to different temperature niches appeared as the main plausible driver explaining the co-exclusion between species A and species B & C (**Figure 2B**). The co-excluding species had different temperature preferences, with species B and C featuring a weighted mean temperature of 27.6 °C, while species A had a weighted mean temperature of 22.1 °C. Both values were significantly different from chance. In contrast, species E did not show any preference associated to temperature in our sample-set covering the tropical and subtropical ocean. A fraction of the taxa positively linked to MAST-4 species showed temperature niche preferences that were coherent with those of species A, B and C (**Figure 2B**; **Supplementary table S5)**. For example, taxa positively associated to species A displayed an average weighted mean temperature of 22.0 °C, while taxa positively associated to MAST-4B/C displayed an average weighted mean temperature of ~26 °C. Both values differed when compared against the average unweighted mean temperature of the entire dataset: ~24°C. Note that detected associations reflecting only environmental preference were removed from the network, therefore remaining positive associations between microbes that prefer similar environmental conditions (e.g. temperature) indicate cases where the links between microbes could not be explained by their comparable environmental preferences. Overall, water temperature explained up to 35% of the variance in the distribution of MAST-4 species (ADONIS, p<0.05).

### Comparative genomics of MAST-4 species

A total of 69 single-cell genomes from MAST-4A (n = 23), MAST-4B (n = 9), MAST-4C (n = 20) and MAST-4E (n = 17) were analyzed. All MAST-4E cells were isolated from the same TARA Oceans station (station 23) at the same depth (DCM) (**Supplementary table S1**). The other MAST-4 single-cells were isolated from different TARA Oceans stations located in either the Indian Ocean or in the Adriatic Sea. These cells originated also from different depths, including Surface or the Deep Chlorophyll Maximum. Based on 18S rRNA-gene similarity, genome tetranucleotide composition and average nucleotide identity, cells of MAST-4A/B/C/E were independently co-assembled (25). The two largest co-assemblies were MAST-4A (47.4 Mb) and MAST-4C (47.8 Mb), which contrasted in terms of size to MAST-4B (29 Mb) and MAST-4E (30.7 Mb). Accordingly, species A and C featured more predicted genes (15,508 and 16,260 respectively) than species B and E (10,019 and 9,042 respectively).

MAST-4 multigene phylogenies based on 30 conserved single-copy predicted proteins (**Supplementary table S3**) as well as genome similarity based on Average Amino acid Identity (AAI) agreed with known phylogenetic relationships based on ribosomal RNA-gene sequences (6) (**Figure 3**). This supports our co-assembly and gene prediction strategy, suggesting also a substantial amount of evolutionary divergence between MAST-4 species A/B/C/E. To assess whether predicted MAST-4 genes have been previously recovered, we compared their nucleotide sequences against the Marine Atlas of Tara Oceans Unigenes (MATOU) (30), a gene catalog of expressed eukaryotic genes clustered at 95% identity into unique genes (unigenes). The number of MAST-4 genes detected in MATOU was variable, with species A featuring ~25%, species B ~20%, species C ~33% and species E ~13% of their predicted genes (**Supplementary table S6**). Not a single MATOU unigene was shared between all four species, while 81.9% of the MATOU unigenes were uniquely associated to one MAST-4 species (**Figure S1**). This supports our previous results indicating a substantial genome differentiation within MAST-4.

**Figure 3.**
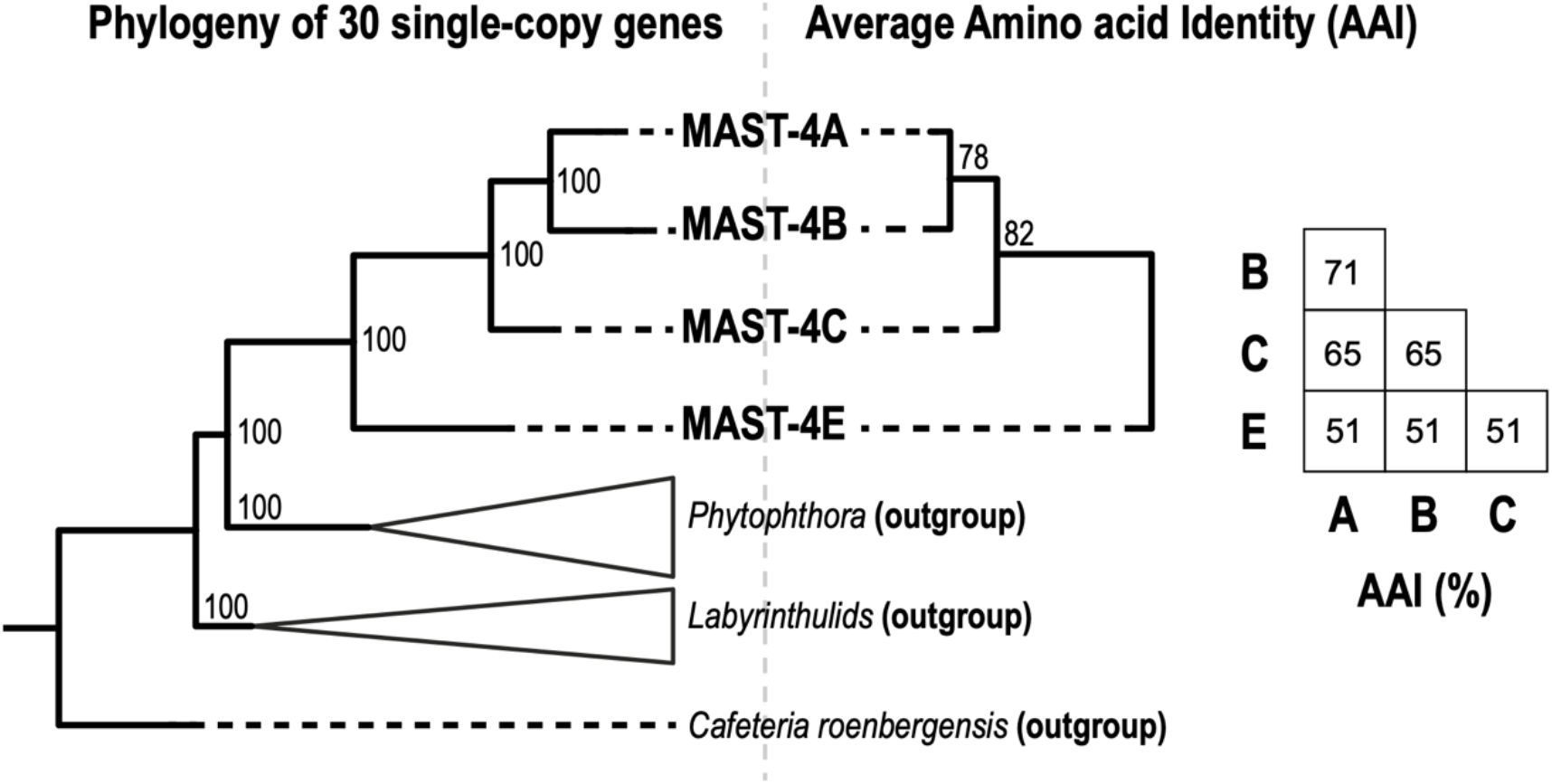
Evolutionary divergence between the studied MAST-4. Left-hand side: MAST-4 species phylogeny based on 30 single-copy protein genes from the BUSCO v3 database (89) that were identified in the co-assemblies (see Methods; **Supplementary Table S3**). Right-hand side: Clustering of MAST-4 co-assembled genomes and bootstrap support based on the Average Amino acid Identity (AAI) between predicted homolog genes. AAI values (%) between MAST-4 species are shown in the matrix.

Predicted amino-acid sequences were functionally annotated using the databases eggNOG, KEGG and CAZy. eggNOG allowed annotating ~75% of the genes from the four species, while ~31% were annotated with KEGG and ~3% with CAZy (**Supplementary table S6**). Considering that eggNOG includes environmental sequences, some with unknown functions, while KEGG and CAZy are based on model or cultured organisms and annotated genes, these differences are not surprising. According to the broad eggNOG functional categories, MAST-4 species shared similar functional profiles (**Figure 4A**). Yet, about half of the eggNOG hits had no function associated, as the reference sequences were environmental. Nevertheless, the existence of these hits further supports our co-assembly and gene prediction approach. The most represented categories with known functions were *‘Posttranslational modification, protein turnover, chaperones’* and *‘Signal transduction mechanisms’*, which group important genes for the proper function of the cell, along with *‘Intracellular trafficking, secretion and vesicular transport’* and *‘Carbohydrate transport and metabolism’*, which include pathways related to food ingestion and degradation (lysosomal reactions). Similarly, KEGG functional categories with the largest number of MAST-4 genes were ‘*Global Metabolism*’, ‘*Signal Transduction*’ and ‘*Transport and Catabolism*’ (**Figure S2**). The first two comprise broad housekeeping functions and pathways, while the third covers vesicular processes such as endo- and phagocytosis. As expected, the potential for grazing is represented in all four MAST-4 genomes

**Figure 4.**
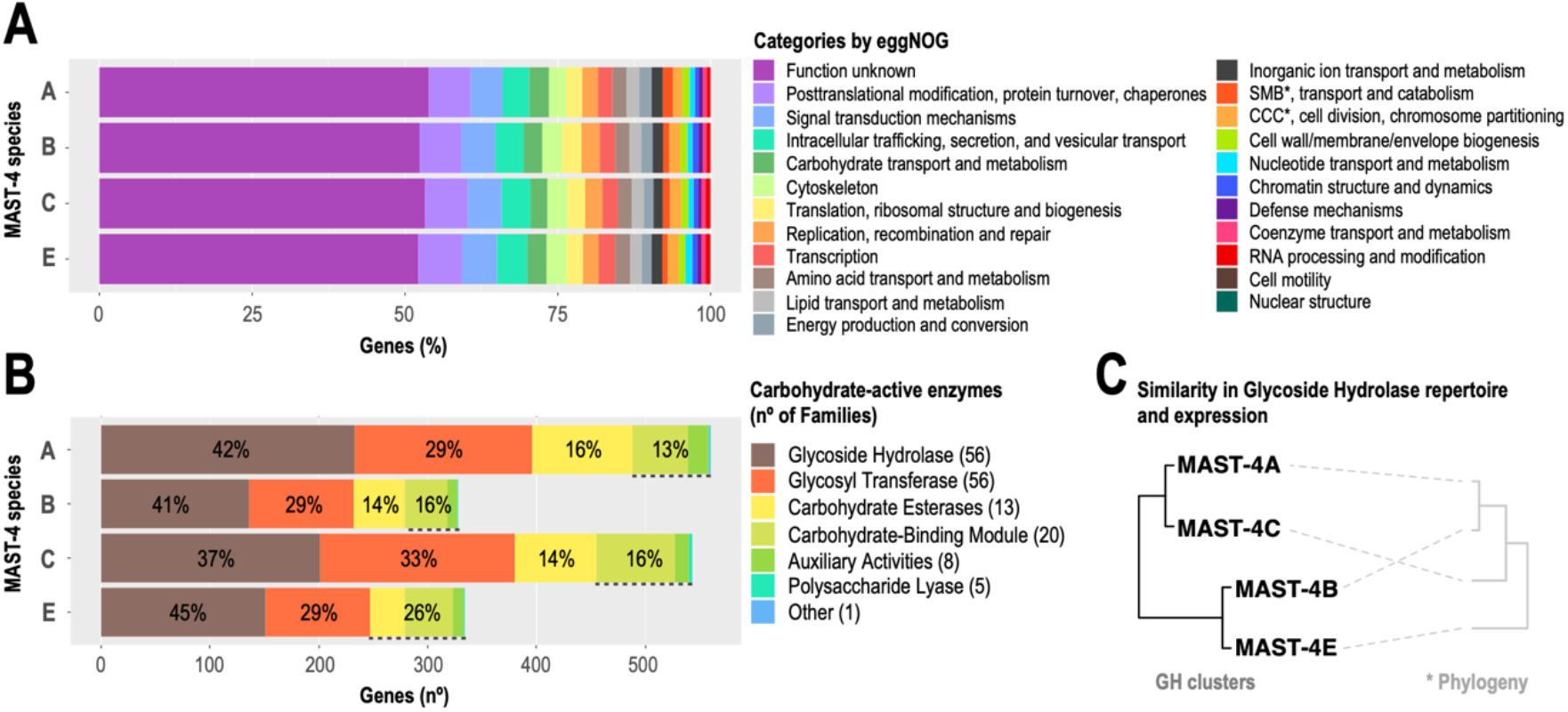
Functional profile of MAST-4 genes according to eggNOG and CAZy. Total MAST-4 genes analyzed were 15,508, 10,019, 16,260 and 9,042 for species A, B, C and E respectively. **Panel A)** eggNOG annotations indicated as percentage of genes falling into functional categories. *SMB – Secondary Metabolites Biosynthesis, CCC – Cell Cycle Control. **Panel B)** Number of MAST-4 genes within CAZy categories and the corresponding percentage; NB: the absolute number of genes is shown between parenthesis. The number of families considered within each CAZy category is indicated in the legend between parenthesis. **Panel C)** Clustering of MAST-4 species using Manhattan distances based on either their Glycoside Hydrolase (GH) composition or the GH expression (in TPM) results in the same clustering pattern. Note that MAST-4C and A are more similar in their GH content than E and B, which are more similar between themselves. **\*** A schematic representation of the phylogeny of the studied MAST-4 is shown for comparison purposes (see Figure 3 for more details).

The CAZy group with the largest number of genes in MAST-4 species was the Glycoside Hydrolases (GHs) (**Figure 4B**). We have analyzed the GH composition of MAST-4, given that different GH repertoires in species could be linked to different capacities to degrade prey bacteria or microalgae (16, 31). Most GH families were found in all MAST-4 species, but some were specific or missing in particular species (e.g. GH23 specific to MAST-4B or GH22 missing in MAST-4E) [**Supplementary table S7**]. Clustering of MAST-4 species based on GH similarity generated two groups, species A – C and B – E (**Figure 4C**). Thus, MAST-4 genomes with contrasting geographic distributions (**Figure 1**) and potential ecological interactions (**Figure 2A**) were clustered together based on GH composition.

### Global expression of MAST-4 Glycoside Hydrolases

Glycoside Hydrolases (GHs) are most likely involved in MAST-4 phagocytosis as parts of the machinery to digest food. We used metatranscriptomic and metagenomic data from the expedition *TARA Oceans* to assess the expression and abundance of MAST-4’s GH genes in the surface global ocean (**Figure 5A**). We found that there was no obvious relationship between GH gene-abundance and expression over the surface global ocean, indicating that differences in gene expression most likely represent up- or down-regulation of GH genes (**Figure 5B**; see also **Figure 5C** and **Figure S3B**). MAST-4’s GH gene expression was highly heterogeneous in the surface global ocean (**Figure 5C**). The GH families with the highest expression were the lysozyme families GH22 and GH24, in charge of degrading the peptidoglycan in the bacterial cell wall (23, 32), as well as the chitinase family GH19, involved in the degradation of chitin (present in particulate detritus, crustaceans and several other organisms in the ocean) [**Figure 5C**]. Interestingly, the South Pacific displayed low or absent GH expression in all MAST-4 species, despite GH gene abundances were similar to those found in other regions displaying higher expression (**Supplementary Figure S3B**). We found also clear differences in expression between species: for example, while species’ A GHs were widely expressed in several regions, those GHs from species’ E were expressed only in specific samples, in particular in the North and South Atlantic. GH genes from species B and C were either not detected or had low expression in the South Atlantic samples, in contrast to specific GH genes from species A and E in the same region (**Figure 5C**). In turn, specific GHs from species B and C had higher expression than A and E in the Indian Ocean.

**Figure 5.**
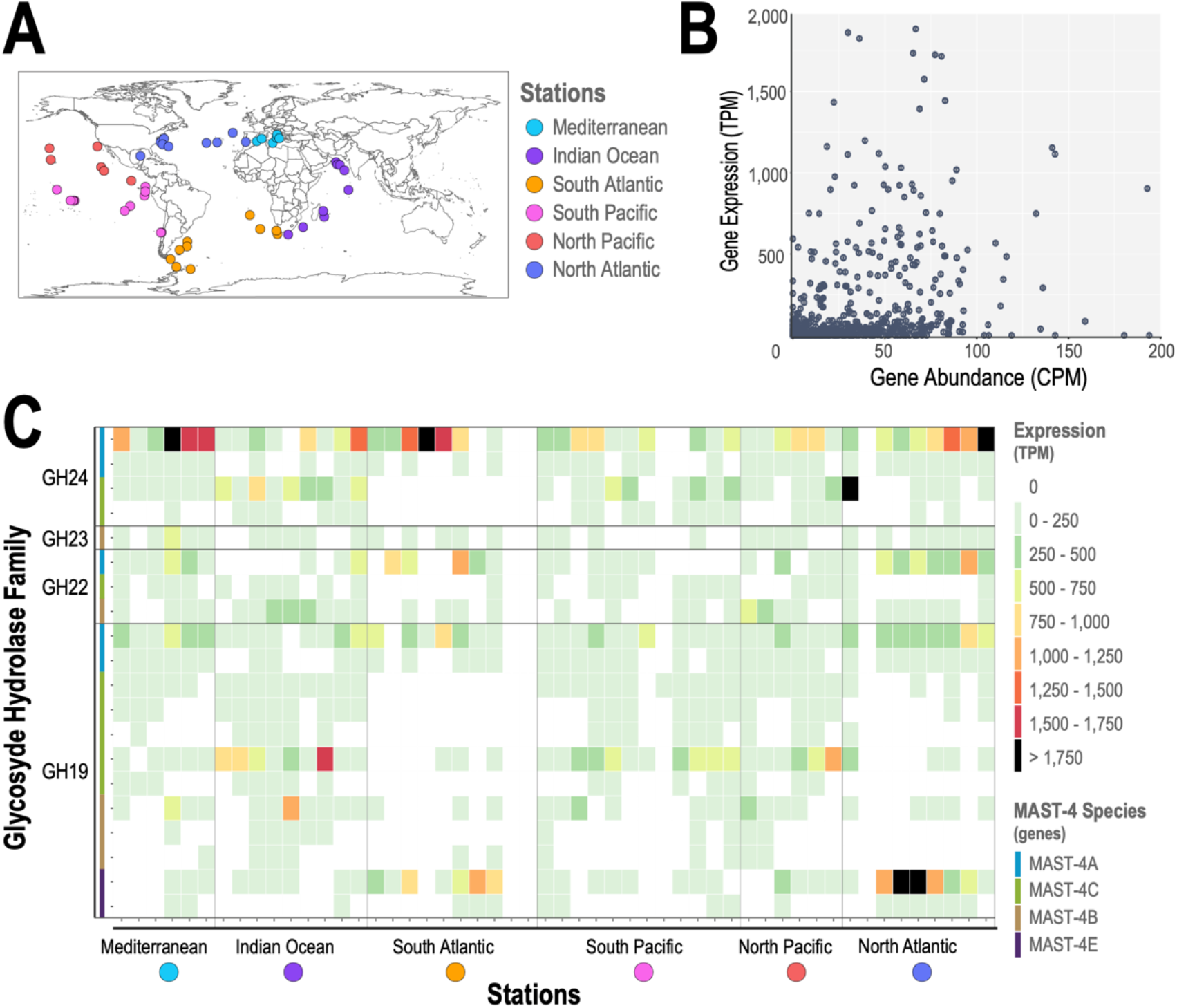
Expression and abundance of GHs in MAST-4A/B/C/E in the upper global ocean. **Panel A)** Geographic location of the metagenomic and metatranscriptomic samples from *Tara Oceans*. **Panel B)** Gene abundance vs. expression using normalized data for each gene and station. Note that the axes have different but proportional ranges of values. **Panel C)** Heatmap of the Glycoside Hydrolase families in MAST-4 that had the highest expression. Samples are in the X-axis, grouped by ocean region and ordered following the expedition’s trajectory. Genes in the Y-axis are organized by family and each species is indicated with a color. GH22, GH23 and GH24 are families of lysozymes and GH19 is a family of chitinases that can also act as lysozyme in some organisms.

Differences in abundance and expression were also found in GH genes belonging to the same family and within the same MAST-4 species. For example, species A had two genes belonging to the GH24 family; one gene (631 bp) was more expressed than the other (1,465 bp), despite gene abundances were similar across all samples (**Figure 5C**; **Supplementary Figure S3B**). These two genes shared 29.5% similarity at the amino acid level based on 73% coverage (153 amino acids) of the shorter gene. A similar pattern was observed in the two GH24 genes in MAST-4C: the shorter was more expressed than the longer one (622 vs. 1,198 bp). In fact, the short and long GH24 genes from species A and C are homologs respectively: the short homologs have 79.4% identity (94% coverage) while the long homologs have 56.4% identity (87% coverage). In general, MAST-4 species with more than one gene belonging to the same GH family tended to express one particular variant over the others. One plausible explanation is that the under-expressed GHs are gene duplications. GH genes often undergo duplication, and thus several copies can be present in the form of paralogs (33–35). After gene duplication, a redundant copy is generated and freed from selective pressure, allowing it to accumulate mutations (36) and potentially lead to new functions (37, 38).

### Detecting positive selection acting on MAST-4 genes

We analyzed whether there is evidence of positive selection leading to niche adaptation in the different MAST-4 species. For that, we analyzed non-synonymous vs. synonymous substitutions (*dN*/*dS*) in selected homologous genes in MAST-4A/B/C/E. Synonymous substitution rates (*dS*) [i.e. they do not produce changes in amino-acids] are generally slow and depend on the species, while non-synonymous rates (*dN*) are even slower due to purifying selection. The action of positive selection is expected to increase *dN*, without affecting *dS*. Normally, the ratio *dN*/*dS* is used to test hypotheses related to the action of selection on protein-coding genes (39). Hence, *dN*/*dS* >1 indicates that substitutions generating changes in amino-acids are higher than substitutions that do not, suggesting the action of diversifying (i.e. positive) selection. In contrast, *dN*/*dS* <1 suggests purifying evolution (i.e. most mutations do not generate changes in amino-acids), while *dN*/*dS* =1 points to neutral evolution.

A total of 692 alignments (homologous groups) were used for testing positive selection on both branch (whole sequence phylogeny) and codon analyses (gene site-specific) (40, 41) (**Supplementary table S8**; see Methods). Overall, 60 gene alignments (8.7%) indicated positive selection in branch analyses, of which 57 alignments displayed selection in one branch and 3 in two branches (60 alignments, 63 total branches selected). MAST-4A and B appeared to be the most selected branches, 22 (34.9%) and 25 (39.7%) times respectively, while MAST-4C and E had a low number of selected branches, 8 (12.7%) and 4 (6.3%) times respectively. In codon analyses, 478 gene alignments (69.1%) displayed positive selection in one or more positions, ranging from 1 to 15 positively selected codons per alignment. In Glycoside Hydrolases (GH), a key part of the predatory machinery of the MAST-4, 1 alignment (0.14%) showed positive selection in branch analyses for family GH74 while 14 alignments (2%) displayed positive selection in codon analyses that included GH3, GH13, GH16, GH19, GH28, GH30, G74, GH78, GH79 and GH99 (**Supplementary table S8**). Of all of them, only GH19 belongs to one of the most expressed families according to the metatranscriptomic analyses. Overall, these analyses suggest that adaptive evolution promoted the diversification of MAST-4 into species A, B, C & E, or at least that it promoted the diversification of these genes.

## DISCUSSION

Currents, waves and wind promote the dispersal of plankton in the surface ocean. Given their typically large populations and small organismal sizes, microbial plankton is expected to be widely distributed in the upper ocean. This is particularly relevant for the MAST-4 group, which features a moderate abundance (about 50 cells ml^−1^ in surface waters and ~10% of the heterotrophic flagellates (42)) and minute size. Such characteristics in combination would guarantee dispersal and widespread distributions (43), decreasing the potential effects of dispersal limitation (44). These characteristics would also promote a coupling between environmental heterogeneity (selection) and species distributions (45). Thus, we expected that MAST-4 distributions would reflect, to a certain extent, the abiotic and biotic conditions in the ocean. This is coherent with previous findings indicating that a) temperature is an important environmental variable driving MAST-4 distributions, and b) that dispersal limitation does not seem to affect the distributions of MAST-4 species (17). We expanded our previous knowledge by determining the temperature distribution of species A, B, and C. Specifically, we show that species B and C occur in warmer temperatures (weighted mean = 27.6 °C), while species A is present in lower temperatures (weighted mean = 22.1 °C). In contrast, we did not find evidence that the distribution of species E was affected by temperature in the tropical and subtropical ocean. This is coherent with the preference of species E for cold waters (17).

Even though temperature is a key variable structuring the global ocean microbiota, including MAST-4 (46–49), biotic variables could also affect the distributions of MAST-4 species. We found that the number of associations between MAST-4 OTUs and bacterial OTUs was low. Actually, most associations were not considered as they were either weak (low correlation) or they just represented similar or different environmental preference (mainly temperature) between MAST-4 and bacterial OTUs. Altogether, this suggests that MAST-4 abundance and occurrence is weakly coupled to bacterial distributions and abundance in the upper ocean, which agrees with previous studies where temperature appeared as the main driver of heterotrophic flagellates abundances (42). We detected a substantial number of taxa that were positively associated to either MAST-4B/C or MAST-4A but not to both. Even though associated taxa tended to reflect the temperature preference of the species to which they were associated (B/C or A), their association to different MAST-4 cannot be simply explained by similar niche temperature, since we also detected associations to OTUs without a significant temperature preference. The vast majority of associations were between species A or B/C with other picoeukaryotes, such as Syndiniales’ Dino-group-I and II, which are known parasites (50), or MAST-3 and MAST-7, which are flagellates as well (6). These associations could either manifest a similar preference for an environmental variable different than temperature that covaries with MAST-4 distributions or reflect real ecological interactions. For instance, there is evidence of MAST-4A having a predator-prey relationship with *Synechococcus* (51) and possibly with SAR11 (12), which was not only reflected in our networks from the *Malaspina* expedition, but also in previous studies from the *Tara Oceans* expedition (52). Results from *TARA Oceans* reported other taxa associated to MAST-4A that were corroborated by our results (MOCH-2, Chrysophyceae, MALVs, MAST-7). However, whether or not these associations reflect true ecological interactions needs to be proved with further experiments. Altogether, we did not find evidence that interspecific biotic interactions represent an important driver of MAST-4 biogeography.

Our results suggest that adaptation to different temperature niches and intraspecific biotic interactions (competition) are likely the main drivers determining MAST-4 biogeography. If so, differential adaptation should likely be reflected in the genomes of the MAST-4 species. Our analyses indicated that MAST-4 species differ in genome size: two bigger genomes (MAST-4A and C) with partial genome size of ~47 Mb and ~80% completeness and two smaller genomes (MAST-4B and E) of ~30 Mb and ~70% completeness, which correspond to ~59 and ~42 Mb full estimated genomes respectively. The observed differences in genome size need to be considered with care, as they may be reflecting incomplete genome assemblies. Nevertheless, our estimates of genome size were similar to those of *Cafeteria roenbergensis* (~40 Mb) (53), an heterotrophic flagellate in the same cell-size range of MAST-4, and other Stramenopile genomes, for example the diatom *Thalassiosira pseudonana* (~34.5 Mb) (54) or specific *Phytophthora* species (*P. plurivora*, *P. multivora*, *P. kernoviae* and *P. agathidicida* with 41, 40, 43 and 37 Mb respectively) (55). This suggests that our partial genomes are likely large enough to be representative of the studied MAST-4 species. We found that differences in MAST-4 genome size were mirrored by the number of predicted genes in each species, which ranged between 9,042 and 16,260, even though larger genomes in eukaryotes do not always imply a greater number of genes (56). These differences in gene content between species may to some extent be linked to niche adaptation. Overall, none of the studied MAST-4 displayed any loss or gain of broad functional categories when compared to each other. In fact, they were similar in terms of proportion of genes that belong to each functional trait, suggesting that MAST-4 metabolisms are broadly comparable, which agrees with other reported results in MAST-4 species A/C and E (16). Among the most represented functional categories in the MAST-4 genomes were those involved in phagocytosis. For instance, eggNOG’s *‘vesicular and carbon transport’*, along with KEGG’s *‘transport and catabolism’*, includes pathways for *‘Endocytosis’*, *‘Phagosome’*, *‘Lysosome’*, *‘Peroxisome’* and *‘Autophagy (animal and yeast)’*, all related to vesicular forms of transport. Thus, MAST-4’s lifestyle as marine grazers (5, 57) appears to be reflected by their broad genomic functions. Yet, homologs among species were very different at the DNA or amino acid level. In particular, when comparing our gene predictions against the *Tara Oceans* gene catalog of marine eukaryotic genes (MATOU) (30, 58), the vast majority of BLAST hits were unique to one MAST-4 species. In fact, we did not find a single gene in MATOU that was present in all four species, which manifests the intraspecific differences in MAST-4 in terms of genome-sequence composition. The substantial differentiation between homologs was reflected by the AAI and phylogenomic results as well (**Figure S1**), which altogether indicate that MAST-4 experienced substantial evolutionary diversification.

MAST-4 is not exclusively bacterivorous and can feed on other small organism, for example *Micromonas pusilla* and *Ostreococcus sp*. (5), and perhaps complement their diets with non-infective viruses (59). A comparable diet has been observed in other heterotrophic flagellates (60). Such plethora of food items, which vary in quality and quantity, most likely require different machineries that are capable to digest them (16, 31), in particular different carbohydrate active enzymes. For example, studies in Fungi have shown how the number and composition of CAZymes may determine the degradation capacity of different plant biomass sources (61). Here, we analyzed the Glycoside Hydrolases (GHs), one of the most efficient known catalysts of organic substances in living organisms (62), and likely important for MAST-4’s heterotrophic lifestyle. GHs genes accounted on average for 3% of the predicted genes in each MAST-4. Most of the GH gene families were found in the four species, but some were either exclusive of a single species or missing in others, which may be due to genome incompleteness. In fact, similar patterns have been reported before, not only in a reduced number of MAST-4 species (16), but also in the fungal genus *Saccharomyces* (63), where the set of GH genes differs even in strains of the same species. Site (codon) analyses suggested positive selection within the GH19 gene family in MAST-4. Similarly, other GH families that are not lysozyme-like, such as GH3, GH30 or GH74 appeared to have experienced positive selection as well, even though they were not as much expressed in the global ocean as the lysozyme. Altogether, this suggests the action of adaptive evolution in the machinery that MAST-4 uses to digest food, and may reflect adaptations to the degradation of different compounds or prey.

The four MAST-4 species formed two groups based on GH similarity (number of genes per family). One group included species A and C and the other species B and E. Interestingly, species A and C, with similar GH repertoires, showed spatial co-exclusion in the upper global ocean, while species C and B, with different GH repertoires, displayed co-occurrence (**Figure 1**). These geographic distributions suggest that niche adaptation associated to different temperatures allowed MAST-4 A and C to keep similar GH repertoires, while species adapted to similar temperatures that co-occur (C and B) were exposed to divergent selection diversifying their diets in order to avoid competition, which is reflected in their different GHs (31). We found that species A and B/C have different niche temperatures (A= 22.1 °C and B/C= 27.6°C). Given that species C and B have almost identical temperature niches, it could have been expected a similar temperature niche for MAST-4B’s closest relative, MAST-4A, considering that temperature niche can be a phylogenetically conserved trait in specific microbes (64, 65). Yet, species A displayed a temperature niche 5 °C lower, suggesting that selection has promoted its adaptation to lower temperatures perhaps to avoid competition with species C, or that species C is a superior competitor an excludes species A from warmer waters. Phylogenetically, it would have been expected that species B had a similar GH repertoire to species A and C, which are the closest evolutionary relatives, but instead it was closest to E, suggesting that evolution promoted the divergence of MAST-4B’s GH content. Altogether, considering the temperature distributions of the studied MAST-4 species, a hypothetical evolutionary scenario unfolds related to temperature adaptation. Here, the more ancestral species E would have adapted to cold temperatures (17), while species C adapted to tropical waters. Then, the two most evolutionary derived species, A and B, diverged their niches to avoid competition with C: species A adapted to colder subtropical and temperate waters, while species B stayed in the tropics by changing its niche via diet modification, and this is reflected in its GH composition. Our *dN*/*dS* results using homologous proteins of the four MAST-4 species are coherent with the previous evolutionary scenario, by indicating that MAST-4A and MAST-4B appear to be the two species under selective pressure, as they displayed significant positive selection in 75% of the total alignments with branch selection.

We also analyzed MAST-4’s GH distribution and expression in the surface global ocean, as this may shed light on whether species with similar GH composition express similar or different genes when they co-occur, possibly indicating prey preference depending on the presence-absence of competitors. We found that the different species displayed a large heterogeneity in their expression patterns. The tropical species that co-occurred the most, C and B, showed dissimilar expression patterns, with some genes being highly expressed only in one species, which is coherent with their difference in GH composition as well as with a scenario proposing different food preference. Furthermore, species C and B showed differences in expression over specific ocean regions, suggesting that despite their co-occurrence, their GH activity is modulated differently. In turn, the co-excluding species A and C, which display the most similar GH composition, appeared to express different GHs over the upper global ocean, suggesting that they regulate GH expression or that GH expression is affected by the different temperatures in which these species occur. Overall, our evidence suggests that species A, B and C regulate GH genes differently, even though some differences in GH expression only reflect the presence or absence of MAST-4 species in specific ocean regions.

Altogether, our results suggest that the evolutionary diversification of MAST-4 was promoted by divergent adaptive evolution towards different temperature and/or diet niches possibly to avoid competition, and that biotic interactions with other species did not have a major influence in MAST-4 diversification. The previous possibly led to the emergence of the species associated to tropical (MAST-4B and C), subtropical-temperate (MAST-4A) and subpolar-polar (MAST-4E) waters. Furthermore, species B may have diverged in its diet to avoid competition with C and as a consequence, it has a different GH composition from its closest evolutionary relatives, A and C. If future cultures of MAST-4 species are established, the previous scenarios could be tested by determining the temperature range of species growing in isolation or with intra-specific competitors. Our work represents a significant contribution to understand the evolution, diversity, biogeography and function of the smallest predators in the ocean. This knowledge is fundamental to comprehend the base of marine food webs and the biotic and abiotic factors that may affect them, as well as the consequences in upper trophic levels.

## METHODS

### Geographic distribution of MAST-4 species and association patterns

The distribution of MAST-4 species as well as their association patterns were investigated using metabarcoding based on data from Logares et al. (49). This dataset includes surface water samples (3 m depth) from a total of 120 globally-distributed stations located in the tropical and sub-tropical ocean that were sampled as part of the *Malaspina 2010* expedition (27). Samples were obtained with a 20 L Niskin bottle deployed simultaneously to a CTD profiler that measured conductivity, temperature, oxygen, fluorescence and turbidity. About 12 L of seawater were filtered to recover the smallest organismal size-fraction (0.2 - 3 μm; picoplankton). The concentration of inorganic nutrients (NO_3_^−^, NO_2_^−^, PO_4_^3−^, SiO_2_) was included in our analyses (see Logares et al. (49) for details on their measurement).

Both the 18S (V4 region (66)) and 16S (V4-V5 region (67)) rRNA-genes were analyzed. Operational Taxonomic Units (OTUs) were delineated as Amplicon Sequence Variants (ASV) using DADA2 (68) and OTU tables were generated. Amplifications were performed with QIAGEN HotStar Taq master mix (Qigen Inc., Valencia, CA, USA). Amplicon libraries were paired-end sequenced using *Illumina* MiSeq (2 × 250 bp) at the Research and Testing Laboratory facility (see Logares et al. (49) for more details). We trimmed the 18S forward reads at 240 bp and the reverse reads at 180 bp, while for the 16S, forward reads were trimmed at 220 bp and reverse reads at 200 bp. Then, for the 18S, the maxEE was set to 7 and 8 for the forward and reverse reads respectively, while for the 16S, the maxEE was set to 2 for the forward reads and to 4 for the reverse reads. OTUs were assigned taxonomy using the naïve Bayesian classifier method (69) together with the SILVA v132 database (70) as implemented in DADA2. Eukaryotic OTUs were also BLASTed against the Protist Ribosomal Reference database (PR^2^, version 4.11.1 (71)). Streptophyta, Metazoa, nucleomorphs, chloroplasts and mitochondria were removed from OTU tables.

To infer associations between OTUs we used eukaryotic and prokaryotic OTUs with total abundances >100 reads and occurrences >15% of the samples. All abundances were center log ratio (clr) transformed. Associations between OTUs were inferred using Maximal Information Coefficient (MIC) analyses as implemented in MICtools (72), which estimates the total information coefficient TIC_e_ and the maximal information coefficient MIC_e_. TIC_e_ is used to estimate significant relationships, while their strength is calculated with MIC_e_. TIC_e_ null distributions were estimated using 200,000 permutations and the significance level was set to 0.001 as suggested by Weiss et al. (73) MIC_e_ = 0 indicates no association between OTUs, while MIC_e_ = 1 indicates strong association. Environmentally-driven associations between OTUs were detected and removed using EnDED (74), with the methods Interaction Information and Data Processing Inequality. Furthermore, to account for data sparsity and the consequential correlations between zeros in the dataset, we removed associations between OTUs that were not present in ≥ 50% of the samples, i.e. less than half of the samples contained at least one of the two OTUs. We determined the Jaccard index for each association based on the presence of OTUs in the samples (intersection divided by union). We removed associations that featured a Jaccard index below 0.25. Moreover, only associations with MIC_e_ > 0.4 were considered. We used the Pearson and Spearman correlation coefficients to analyze the association type: positive Pearson or Spearman correlation coefficients point to co-occurrences, while negative values point to mutual exclusions. The distribution of OTUs across sea temperatures was explored using the *niche.val* function in the EcolUtils package (75). The abundance-weighted mean temperature was calculated for each OTU and used as an estimate of its temperature niche. We checked whether the obtained abundance-weighted mean temperature for each OTU was significantly different from chance (p<0.05) using a null model with 1,000 randomizations.

### Genome reconstruction using Single Amplified Genomes

Plankton samples were collected during the circumglobal *Tara Oceans* expedition and cryopreserved as described elsewhere (76). Individual picoplankton cells were isolated from water samples and stained with 1x SYBR Green I (Life Technologies Corporation) (25, 77) using a MoFlo (Dako Cytomation Carpinteria, CA, USA) flow cytometer equipped with the CyClone robotic arm for sorting into plates of 384 wells. Cells were lysed and their DNA denatured using cold KOH. The genome from each single cell was amplified using Multiple Displacement Amplification (MDA) based on the Phi29 polymerase (RepliPHI™, Epicentre Biotechnologies, Madison, WI, USA) (12, 78). All single-cell work was performed at the Single Cell Genomics Center (htps://scgc.bigelow.org). The obtained SAGs were taxonomically screened by PCR amplification and Sanger sequencing of the 18S rRNA gene using universal eukaryotic primers. A total of 69 SAGs affiliating to MAST-4 species A/B/C/E were selected for downstream analyses. Each selected MAST-4 SAG was sequenced in 1/8 of a lane using either *Illumina* HiSeq2000 or HiSeq4000 at either the Oregon Health & Science University (USA) or the French National Sequencing Center (Genoscope, France). A total of 424.1 Gb of sequencing data was produced, averaging 6.1 (± 0.22) Gb per SAG. For each SAG, sampling location, depth and date are reported in **Supplementary Table S1**.

Each SAG was *de novo* assembled using SPAdes 3.10 (79) in single cell mode “-sc” with default parameters. Contigs shorter than 1kbp were discarded. Quality control and general assembly statistics were computed with Quast v4.5 (80). Estimation of genome recovery was calculated with BUSCO v3 (Benchmarking Universal Single-Copy Orthologs) (81) using the Eukaryota_odb9 dataset (**Supplementary table S2**). SAGs were also co-assembled in order to increase genome recovery. Only SAGs belonging to putatively the same species were co-assembled. Thus, SAGs had to fulfill three conditions to be co-assembled: *First*, their 18S rRNA-gene amplicon needed to be >99.5% similar. *Second*, their Average Nucleotide Identity (ANI) had to be >95%; ANI was computed using Enveomics (82) with the full-length contigs of all SAGs within each species. *Third*, SAGs had to display an homogeneous composition in Emergent Self-Organizing Maps (ESOM) (83) based on tetranucleotide frequencies. Tetranucleotide frequencies were computed using a 1bp sliding window in fragmented contigs between 2.5 and 5 kbp in size considering both DNA strands, and were subsequently clustered using ESOM. Raw data were normalized using robust estimates of mean and variance (“Robust ZT” option) and trained with the k-Batch algorithm and Euclidean grid distance. If fragments from a given SAG were mixed with those from another SAGs in tetranucleotide ESOM representations, it indicated that their genomic structure was similar. SAGs fulfilling the previous three criteria were considered to belong to the same species and were subsequently co-assembled. Three MAST-4C SAGs (AB536_E17, AB536_F22, AB536_M21) showed more genomic divergence (ANI ~93%) compared to the others but were still included in the final co-assembly because the 18S and tetranucleotide frequencies passed the thresholds.

A total of 69 SAGs belonging to MAST-4 were co-assembled: MAST-4A (23 SAGs), MAST-4B (9 SAGs), MAST-4C (20 SAGs) and MAST-4E (17 SAGs). Prior to co-assembly, reads were digitally-normalized using BBNorm (84), considering a minimum coverage depth of 5x and a maximum target coverage depth of 100x. Normalized reads were co-assembled with SPAdes 3.10 using the single-cell mode (“--sc”) running only the assembly module (“--only-assembler”). To extend contigs, they were re-scaffolded with SSPACE v3 (85). Repetitive regions were masked, along with tRNA sequences, using RepeatMasker (86) and tRNAscan-SE-1.3 (87). Quality and assembly statistics were computed with Quast (80) and are shown in **Supplementary table S2**. Parameters not mentioned were set to default. Co-assembled SAGs were carefully checked for foreign DNA. Based on the premise that sequences from the same species have virtually the same tetra-nucleotide frequencies, a second tetra-nucleotide ESOM map was built for the four MAST-4 co-assemblies with the same parameters as previously described. Contigs that did not cluster together with the majority of contigs from a given SAG co-assembly were removed. Subsequently, co-assembled contigs that were classified as prokaryotic were removed based on the 5-mer profiles using EukRep (88) with mild stringency. Lastly, eukaryotic contigs with extreme GC content values, *i.e*. values outside the range of GC content mean ± 10 % (Standard deviation) in each SAG co-assembly, were removed as well (**Supplementary table S2**). Co-assembled genome completeness was estimated with BUSCO v3 (89). For each co-assembly, protein-coding genes were predicted *de novo* with AUGUSTUS 3.2.3 (90, 91) using the identified BUSCO v3 proteins as training set. Predicted genes were functionally annotated using 1) CAZy database from dbCAN v6 (92) and HMMER 3.1b2 (93) (e-value ≤ 10^−5^), 2) KEGG (Release 2015-10-12; (94, 95)) and 3) eggNOG v4.5 (96), both using BLAST 2.2.28+ and considering hits with >25% identity, >60% query coverage, <10^−5^ e-value and amino acid alignment lengths >200. Gene sequences (nucleotides) were also mapped against the Marine Atlas of Tara Oceans Unigenes (MATOU) Version 1 (20171115)(30) using BLAST 2.2.28+ with the same thresholds as the ones above used for the amino acid sequences, except for the identity threshold, which was increased to 75%, to consider nucleotide sequence variation instead of amino acid.

### Phylogenomics and genome differentiation

We used two approaches to analyze the phylogenetic vs. whole-genome differentiation among MAST-4 species. In the *first* approach, we randomly selected 30 conserved proteins (included in eukaryota_odb9, BUSCO v3) that were identified in all MAST-4 species (**Supplementary table S3**) as well as in other publicly available Stramenopile genomes: *Phytophtora sojae* (NCBI:txid67593), *Phytophtora infestans* (NCBI:txid403677), *Schizochytrium aggregatum* (JGI:Schag1)*, Aurantiochytrium limacinum* (JGI:Aurli1) and *Cafeteria Roenbergensis* (53). Genes were aligned individually with Mafft (97) using the ‘—auto’ mode and concatenated with catfasta2phyml (98). Poorly aligned sequences and regions were removed using trimAl v1.4.rev22 (99) under the “-automated1” mode and default parameters. The phylogenetic tree was built with RAxML version 8.0.0 (100) using the General Time Reversible model with a gamma distributed rate variation among sites (GTR+G). Initial seed was “-p 666”. In addition, we used the automatic bootstrap criterion (-autoMRE) and rapid Bootstrap mode (-f a). The *second* approach consisted in computing the Average Amino-acid Identity (AAI) for each pair of MAST-4 using Enveomics based on the predicted genes (amino acids). Genomes were clustered by similarity using the *pvclust* (101) package in R with “maximum” as distance method.

### Abundance and expression of selected MAST-4 ERGs in the ocean

We investigated the distribution, abundance and expression in the global ocean of selected Ecological Relevant Genes (ERGs), in this case, lysosomal enzymes (glycoside hydrolases). For that, we mapped metagenomic and metatranscriptomic reads from *TARA Oceans* (a total of 52 surface water stations encompassing the 0.8 – 5 μm size fraction (total 104 samples), the range where MAST-4 is found) against predicted genes from each MAST-4 species (**Supplementary table S4**). The mapping was done with BWA (102) and only hits with identity > 95% and an alignment length > 80 bp were considered. Reads that mapped to more than one target were discarded. Gene abundance and expression estimates were normalized by dividing the Reads Per Kilobase (RPK) of each gene [number of mapped reads (counts) / gene length (kbp)] by the Scaling Factor (SF) [Sum of all considered RPKs in a sample / 10^6^]. Hereafter, the abundance of genes and transcripts is expressed as Counts Per Million (CPM) or Transcripts Per Million (TPM) respectively.

### Calculation of dN/dS ratios in homologous genes

Homologous MAST-4 genes were identified using reciprocal protein BLAST (v. 2.2.28+) with the following thresholds: >25% identity, >60% of query coverage, <10^−5^ e-value and an alignment length >200 amino acids. Gene sequences (amino acid) were aligned using Mafft 7.402 with default parameters and then converted into a codon-based nucleotide alignment with Pal2nal (103). Alignments with one or more unknown nucleotides (Ns) were discarded. For each homolog, a nucleotide-based phylogenetic tree was built using RAxML 8.2.12 (100), with the model GTR+CAT, including bootstrap analyses, and a starting seed “-p 12345” as well as the optimization “-d” parameter. Positive selection was tested on each homolog with HyPhy 2.3.14 (104) using aBSREL (branch) (40) and MEME (site) (41) models considering the codon-based nucleotide alignment and the previous phylogenetic tree. Parameters included options for universal code and testing in all branches. A *p-value* of 0.1 (default) was used for the analysis with the MEME model.

## ACKNOWLEDGMENTS

We thank all scientists from the *Malaspina 2010* expedition and the *Tara Oceans 2009-2013* expedition as well as the staff of the Single Cell Genomics Center (Bigelow Laboratory) for the generation of single amplified genomes. Bioinformatics analyses were performed at the MARBITS platform of the Institut de Ciències del Mar (ICM; http://marbits.icm.csic.es) and also on resources provided by UNINETT Sigma2 - the National Infrastructure for High Performance Computing and Data Storage in Norway. We thank Lidia Montiel and Pablo Sánchez for the assistance with bioinformatics. FL was supported by the Spanish National Program FPI 2016 (BES-2016-076317, MICINN, Spain). RL was supported by a Ramón y Cajal fellowship (RYC-2013-12554, MINECO, Spain). This work was supported by the project INTERACTOMICS (CTM2015-69936-P, MINECO, Spain to RL) and MicroEcoSystems (240904, RCN, Norway to RL). IMD and AL were supported by the European Union’s Horizon 2020 research and innovation program under the Marie Skłodowska-Curie grant agreement no. 675752 (Singek: http://www.singek.eu). We thank the CSIC Open Access Publication Support Initiative through the Unit of Information Resources for Research (URICI) for helping to cover publication fees.

