## Supplementary material for "Evolutionary diversification of tiny ocean predators": Figure S

### Supplementary Figures

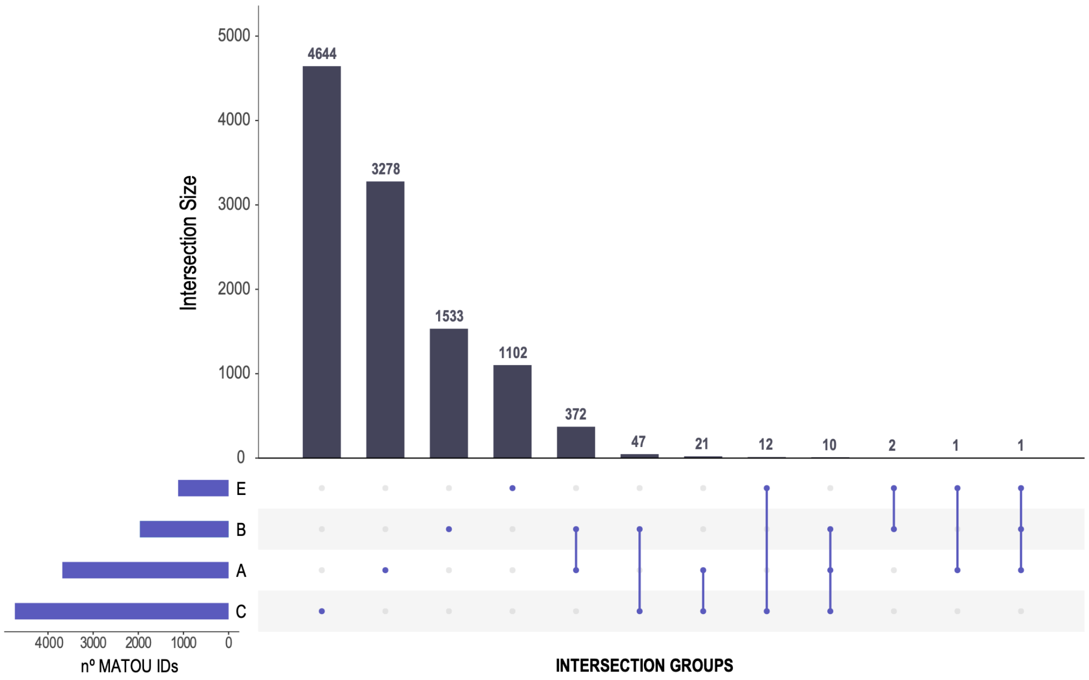


**Figure S1.** Number of Unigenes (i.e. representative genes after clustering genes at 95% identity) from the MATOU database found in MAST-4 along with the number of genes shared by the four species. Note that the different groups are ordered by group size and that the biggest groups are those including only one MAST-4 species, followed by the groups constituted by the combination of two or more species

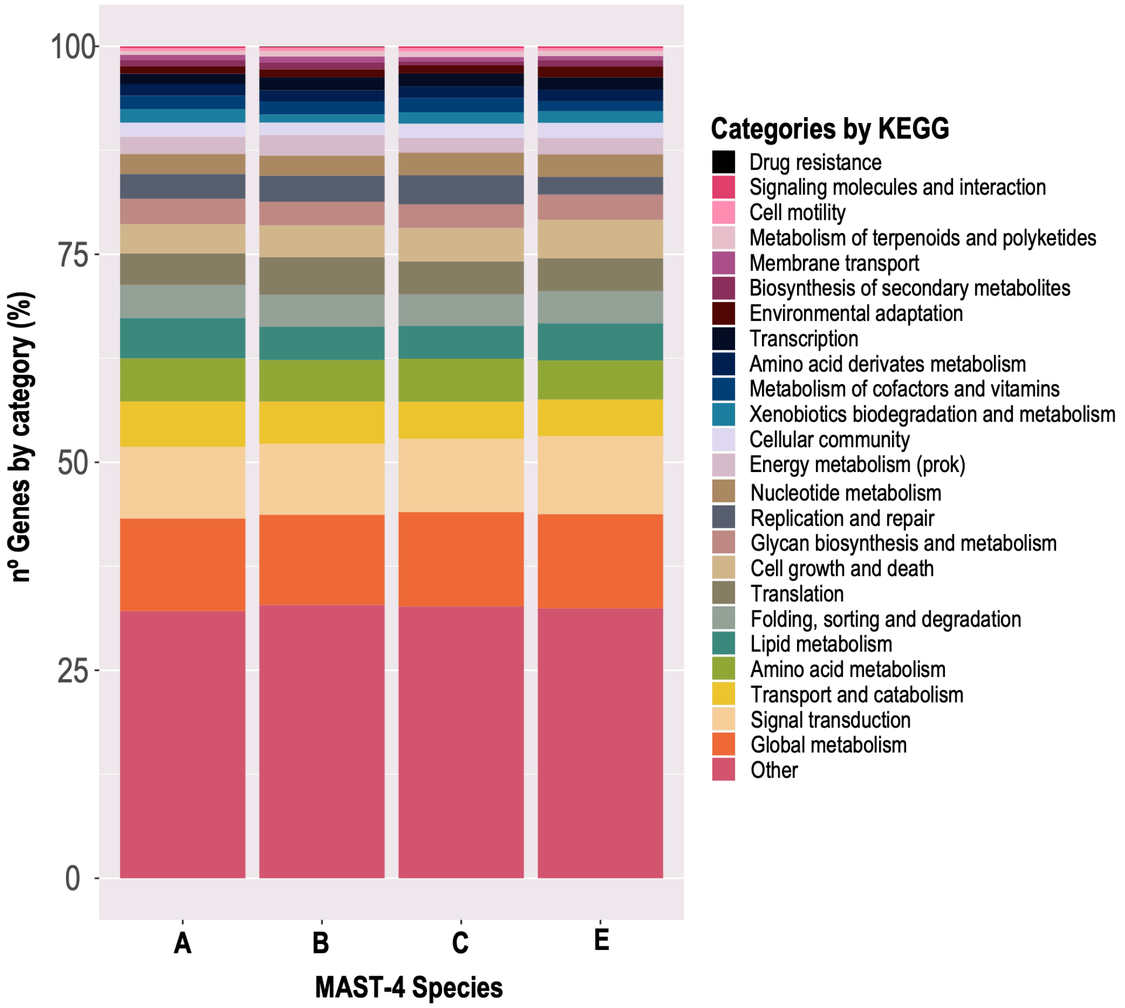


**Figure S2**. Functional profile of MAST-4 genes according to KEGG. KEGG annotations are indicated as percentage of genes falling into functional categories. The category “Other” is an artificial grouping including all the annotations belonging to human related pathways such as ‘Alzheimer’ or ‘Influenza A’.

# A

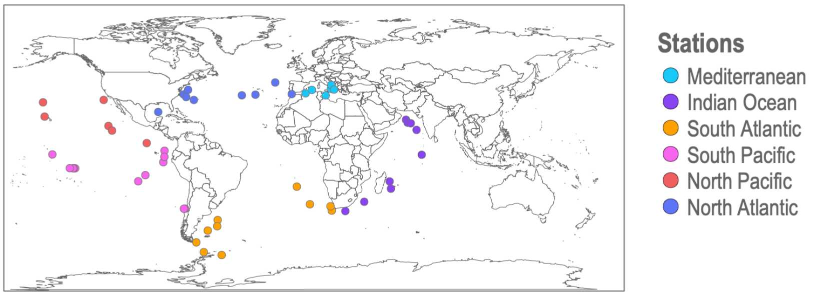


# B

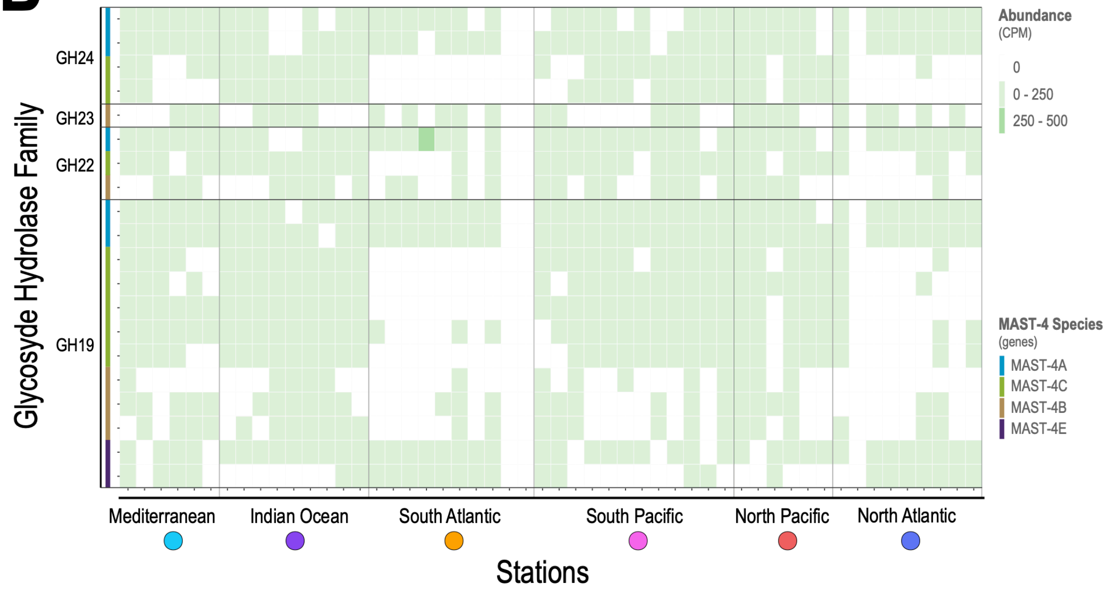


**Figure S3**. Abundance of lysosomal genes in MAST-4A/B/C/E. **Panel A)** Geographic location of metagenomic and metatranscriptomic samples of Tara Oceans. **Panel B)** Heatmap of the Glycoside Hydrolase family abundances in MAST-4 (see their expression in Figure 5C). Samples are in the x-axis grouped by the ocean region and ordered following the expedition's trajectory. Genes in the y-axis are organized by family and each species is indicated with a color. GH22, GH23 and GH24 are families of lysozymes and GH19 is a family of chitinases that can also act as lysozymes in some organisms.
